# The Human STAT2 Coiled-Coil Domain Contains a Degron for Zika Virus Interferon Evasion

**DOI:** 10.1101/2021.10.01.462781

**Authors:** Jean-Patrick Parisien, Jessica J. Lenoir, Gloria Alvarado, Curt M. Horvath

## Abstract

The ability of viruses to evade the host antiviral immune system determines their level of replication fitness, species specificity, and pathogenic potential. Flaviviruses rely on the subversion of innate immune barriers including the type I and type III IFN antiviral systems. Zika virus infection induces the degradation of STAT2, an essential component of the IFN stimulated gene transcription factor, ISGF3. The mechanisms that lead to STAT2 degradation by Zika virus are poorly understood, but it is known to be mediated by the viral NS5 protein that binds to STAT2 and targets it for proteasome-mediated destruction. To better understand how NS5 engages and degrades STAT2, functional analysis of the protein interactions that lead to Zika virus and NS5-dependent STAT2 proteolysis were investigated. Data implicate the STAT2 coiled-coil domain as necessary and sufficient for NS5 interaction and proteasome degradation after Zika virus infection. Molecular dissection reveals that the first two α-helices of the STAT2 coiled-coil contain a specific targeting region for IFN antagonism. These functional interactions provide a more complete understanding of the essential protein-protein interactions needed for Zika virus evasion of the host antiviral response, and identifies new targets for antiviral therapeutic approaches.

**Importance:** Zika virus infection can cause mild fever, rash, and muscle pain, and in rare cases lead to brain or nervous system diseases including Guillain–Barré syndrome. Infections in pregnant women can increase the risk of miscarriage or serious birth defects including brain anomalies and microcephaly. There are no drugs or vaccines for Zika disease. Zika virus is known to break down the host antiviral immune response, and this research project reveals how the virus suppresses interferon signaling, and may reveal therapeutic vulnerabilities.

## Introduction

Interferon (IFN) family cytokines are recognized as fundamental mediators of innate antiviral responses that restrict virus replication and establish essential tissue-specific and species-specific barriers to virus infection (1). Type I IFN includes the single IFNβ and multiple IFNα subtypes that bind to a common receptor, and Type III IFN includes IFNs-λ1-4 that bind to a distinct receptor (2-5). Type I IFN induces the expression of hundreds of antiviral effector genes in nearly all nucleated cell types (6-8), but the Type III IFN system is particularly important for protecting mucosal epithelial cells and establishing barrier functions of tissues. Although these two antiviral IFNs use distinct receptors, their intracellular signaling converges on the formation of ISGF3, a complex of IRF9 along with IFN-activated STAT1 and STAT2 (9). Consequently, these cytokine families activate an overlapping antiviral gene expression program leading to hundreds of IFN-stimulated genes, or ISGs (10-12). The accumulation of ISG products produces an antiviral state that creates a broadly effective barrier that is a hostile environment for virus replication. STAT2 plays an essential role in Type I and Type III IFN responses and is a master regulator of both tonic and acute ISG transcription and antiviral gene regulation (9, 13, 14). For most viruses, the IFN system provides strong selective pressure leading to the evolution of evasion and antagonism strategies (15-17) that can determine their replication efficiency, host range, virulence, and pathogenicity.

Zika virus (ZV) is a member of the Flavivirus family with significant medical consequences for infected patients as well as their unborn offspring. Typically transmitted via *Aedes* mosquitos, ZV typically causes a mild to moderate illness in adults, but can also drive more serious symptoms including Guillain–Barré syndrome. In the developing fetus, ZV is capable of causing severe microcephaly, brain development defects, ocular anomalies, and other features of neurological impairment (18, 19). There are no current anti-Zika therapies or vaccines available, indicating an urgent need for understanding the mechanisms this virus uses to counteract immunity in its human host, and for developing new treatment methods (20).

Like other Flaviviruses, ZV has a single-stranded positive-sense RNA genome that is translated as a single polyprotein, and processed by host and viral proteases to generate the structural proteins (capsid (C), pre-membrane (prM), and envelope (E)), and the non-structural proteins required for host suppression, genome replication, and protein processing (NS1, NS2A, NS2B, NS3, NS4A, NS4B, and NS5). Several Flavivirus NS proteins have been shown to alter host immune recognition or mediate antagonism of host antiviral responses (21). Among these, the NS5 protein has been shown to inhibit IFN signaling by targeting STAT2 (16, 22). Zika virus NS5 protein can target human STAT2 for proteasome-mediated degradation, but the mechanisms leading to STAT2 degradation are poorly understood. The closely-related Dengue virus NS5 protein can target STAT2 for destruction by hijacking the cellular UBR4 protein (23), but ZV-mediated STAT2 degradation apparently does not require UBR4 (24). To gain greater insight into the mechanisms by which ZV NS5 targets the IFN response, the protein interactions that lead to STAT2 proteasome targeting were investigated. Results indicate that the STAT2 coiled-coil (CC) domain is necessary and sufficient for mediating ZV NS5 interaction and initiating degradation via the proteasome after ZV infection. Moreover, the first two α-helices of the STAT2 coiled-coil domain constitute a minimal targeting region for ZV IFN antagonism. These functional interactions provide a more nuanced and detailed understanding of the essential protein-protein interactions needed for ZV to evade the host antiviral response, and they represent potential targets for developing anti-ZV therapeutic approaches.

## Results

### The STAT2 N-terminus mediates Zika virus NS5 interaction

In support of prior investigations of ZV IFN evasion, we observed that infection of cultured primate cells results in loss of STAT2, but has no apparent effect on the closely related STAT1 (Figure 1A). In agreement with this paradigm, expression of FLAG-tagged ZV NS5 by transient transfection co-immunoprecipitates with endogenous STAT2 (Figure 1B) and is able to suppress IFN-stimulated transcription (Figure 1C,D). These results are consistent with previous findings regarding specific STAT2 and IFN targeting by NS5 (16, 22). STAT2 shares the general domain structure of other STAT proteins (25), with an N-terminal domain (ND) adjacent to an extended 4-helix coiled-coil (CC) upstream of the DNA binding (DBD), linker (LD), SH2, and transcriptional activation domains (TAD; see Figure 2A and 2C). To define minimal STAT2 fragments that interact with ZV NS5 and lead to proteasomal degradation, experiments were carried out by expressing ZV NS5 with full length human STAT2 or a partially-overlapping set of STAT2 fragments (Figure 2A). STAT2 proteins containing the first 315 amino acids, representing the STAT2 ND and CC regions, were able to co-precipitate with NS5, but the DNA-binding domain, linker domain, SH2 and TAD were not (Figure 2B). In a corresponding experiment, a complementary set of chimeric STAT1/2 proteins with swapped ND/CC domains were assessed for NS5 association (Figure 2C). Only the chimeric STAT protein containing amino acids 1-315 of STAT2 was able to co-precipitate with ZV NS5 (Figure 2D, N2C1), but the complementary protein with the STAT1 ND and CC was not (Figure 2D, N1C2). These results suggest that the ND and/or CC of STAT2 serve as the target for NS5 recognition.

**Figure 1.**
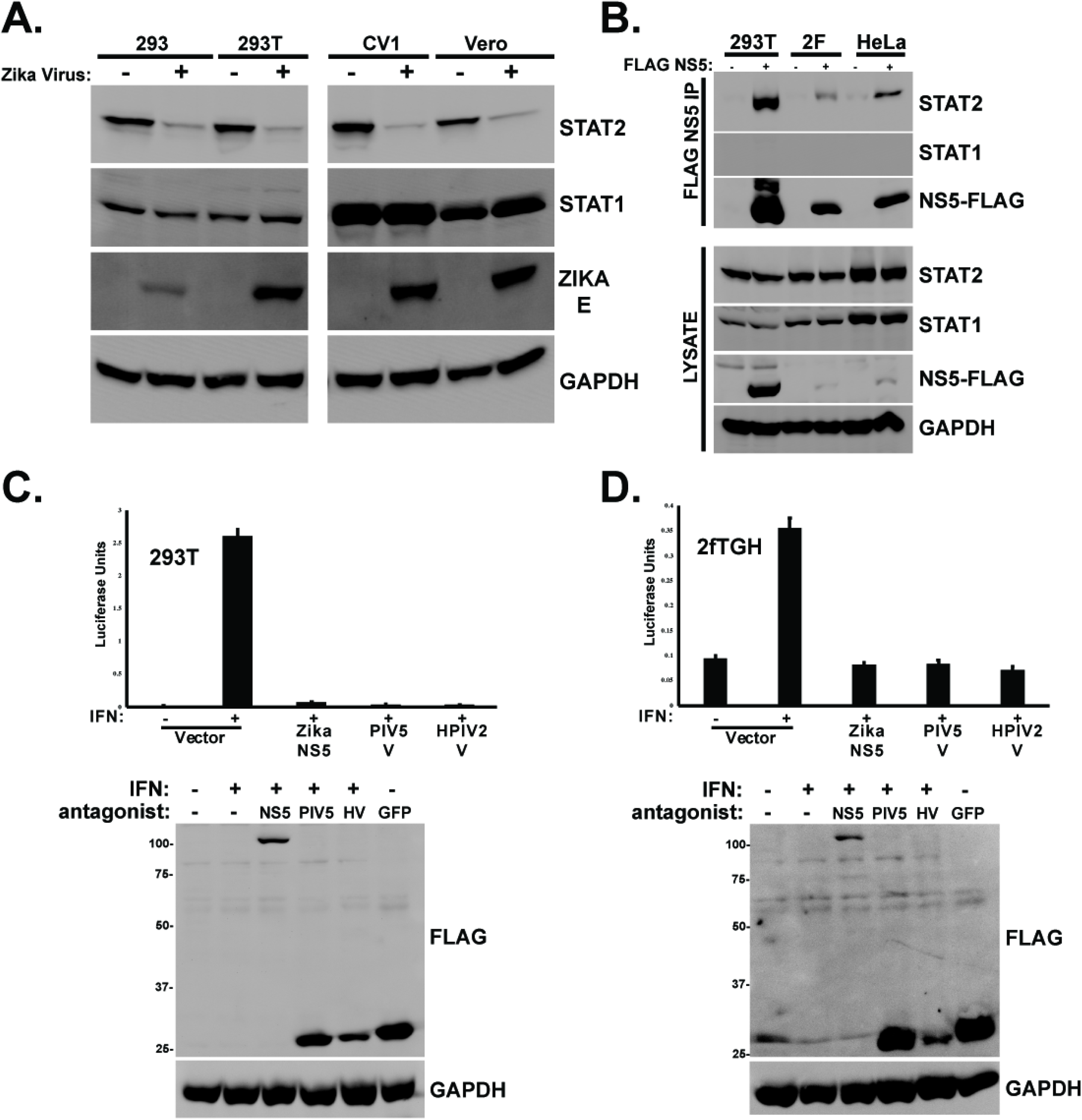
Zika virus STAT2 targeting and degradation requires amino acids 1-315. A.) Infection of primate cells with Zika virus results in STAT2 degradation. Cells were infected with 5 pfu/cell ZV MR766 and incubated 48h prior to lysis and immunoblot with antiserum for STAT2, STAT1, Zika E protein, or GAPDH. B.) Zika virus NS5 co-precipitates with endogenous STAT2. FLAG-tagged NS5 was expressed by plasmid transfection into indicated cell lines, and lysates immunoaffinity purified with FLAG M2 agarose prior to immunoblot with antiserum for STAT2, STAT1, GAPDH, or FLAG peptide. C.) IFN-stimulated gene expression is blocked by ZV NS5. ISRE-luciferase reporter gene was transfected into HEK293T cells with empty expression vector or plasmids encoding NS5 or the V proteins from paramyxoviruses PIV5 or HPIV2, and stimulated with 1000U/ml IFNα for 8h prior to luciferase assay. Data normalized to co-expressed Renilla luciferase; bars represent average of n=3 ±SD. Immunoblot controls below were probed with anti-FLAG antiserum to detect viral IFN antagonist expression (ZV NS5, PIV5 V, and HPIV2 V) as indicated. FLAG GFP included as a negative control and indicator of transfection efficiency. D.). Similar to C., but using human fibrosarcoma 2fTGH cells. Immunoblot controls below were probed with anti-FLAG antiserum to detect viral IFN antagonist expression (ZV NS5, PIV5 V, and HPIV2 V) as indicated. FLAG GFP included as a negative control and indicator of transfection efficiency.

**Figure 2.**
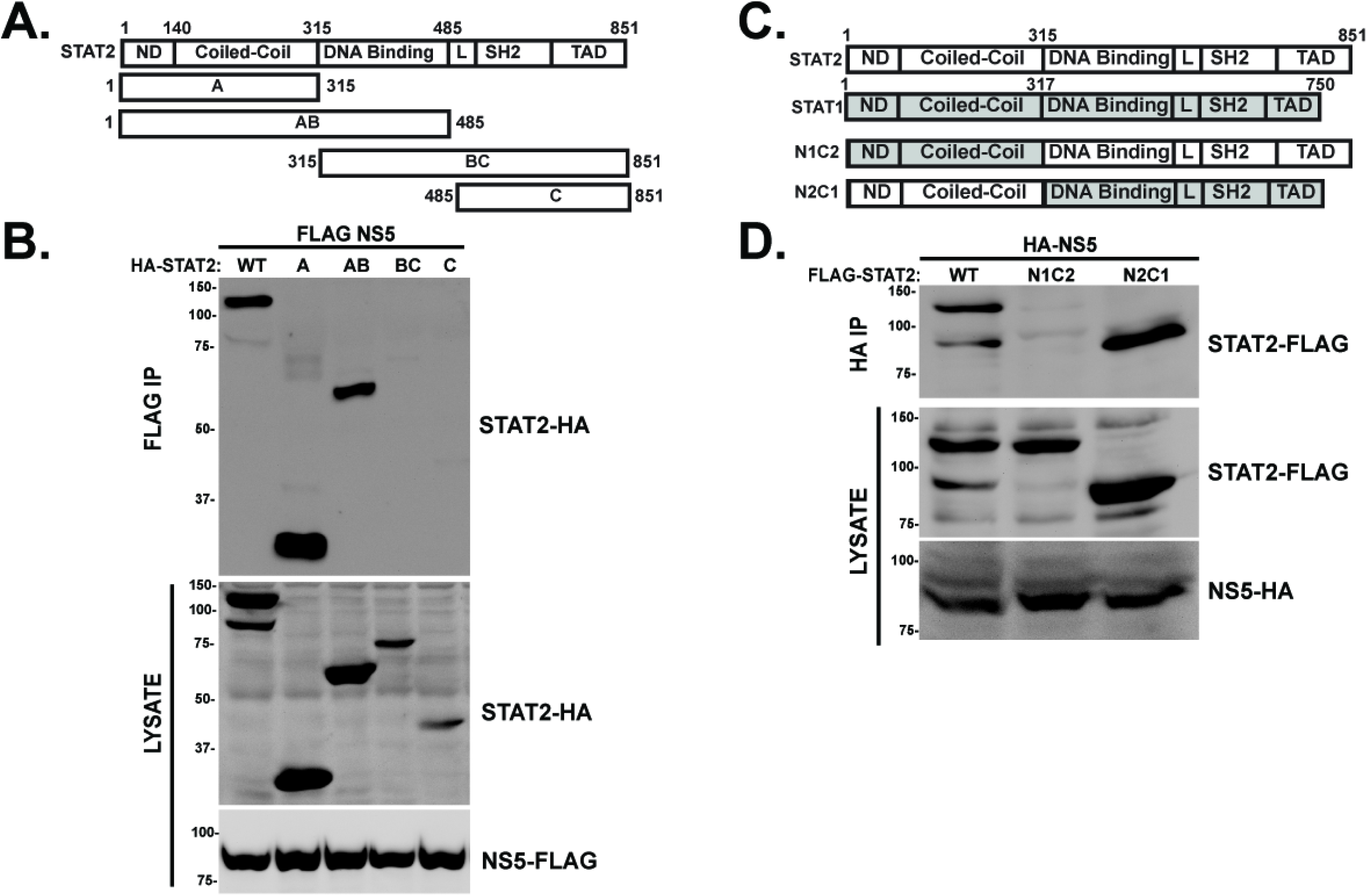
STAT2 N-terminal domains specify recognition by Zika Virus NS5. A.) Diagram of full-length WT STAT2 and STAT2 fragments A, AB, BC, and C with domains and boundaries indicated. B.) STAT2 amino acids 1-315 engage ZV NS5. HA-tagged STAT2 proteins were expressed in HEK 293T cells along with FLAG-tagged NS5, and lysates were immunoaffinity purified with FLAG M2 agarose prior to immunoblot with HA antibody to detect co-precipitating fragments. C.) Diagram of STAT2 (white), STAT1 (grey), and chimeric N1C2 and N2C1, with domain boundaries indicated. D.) STAT2 amino acids 1-315 are transferrable for NS5 recognition. FLAG-tagged STAT2, N1:C2, and N2:C1 proteins were expressed along with HA-tagged NS5, and immunoprecipitated with HA-specific antibody prior to immunoblot with FLAG antiserum to detect co-precipitation.

The STAT protein ND and CC domains are known to mediate protein interactions important for signaling, ISGF3 assembly, and co-factor association, and recent structural studies implicated both domains in ZV NS5 interaction (2, 25-30). To evaluate the contribution of the STAT2 domains toward NS5 recognition and degradation in intact cells, stable cell lines were selected expressing full length STAT2 and its fragments as GFP fusion proteins (Figure 3A). The ability of ZV to degrade the full length, WT STAT2-GFP fusion protein was verified and quantified with flow cytometry. Infection with 5 pfu/cell ZV potently reduced the level of the GFP-STAT2 to 41% by 24 hpi, and to 14% after 48 hpi (Figure 3B). GFP fusions with the combined STAT2 ND and CC (aa 1-315), and the isolated STAT2 ND (1-140) were similarly expressed and detected by immunoblot in uninfected cells. Zika virus infection induced degradation of endogenous STAT2, and GFP fusions to full-length WT STAT2 and the NDCC fragment (Figure 3C). In contrast, neither GFP-STAT1 nor GFP-STAT2-ND fusions were targeted by ZV infection, though the endogenous cellular STAT2 protein remained susceptible to ZV degradation.

**Figure 3.**
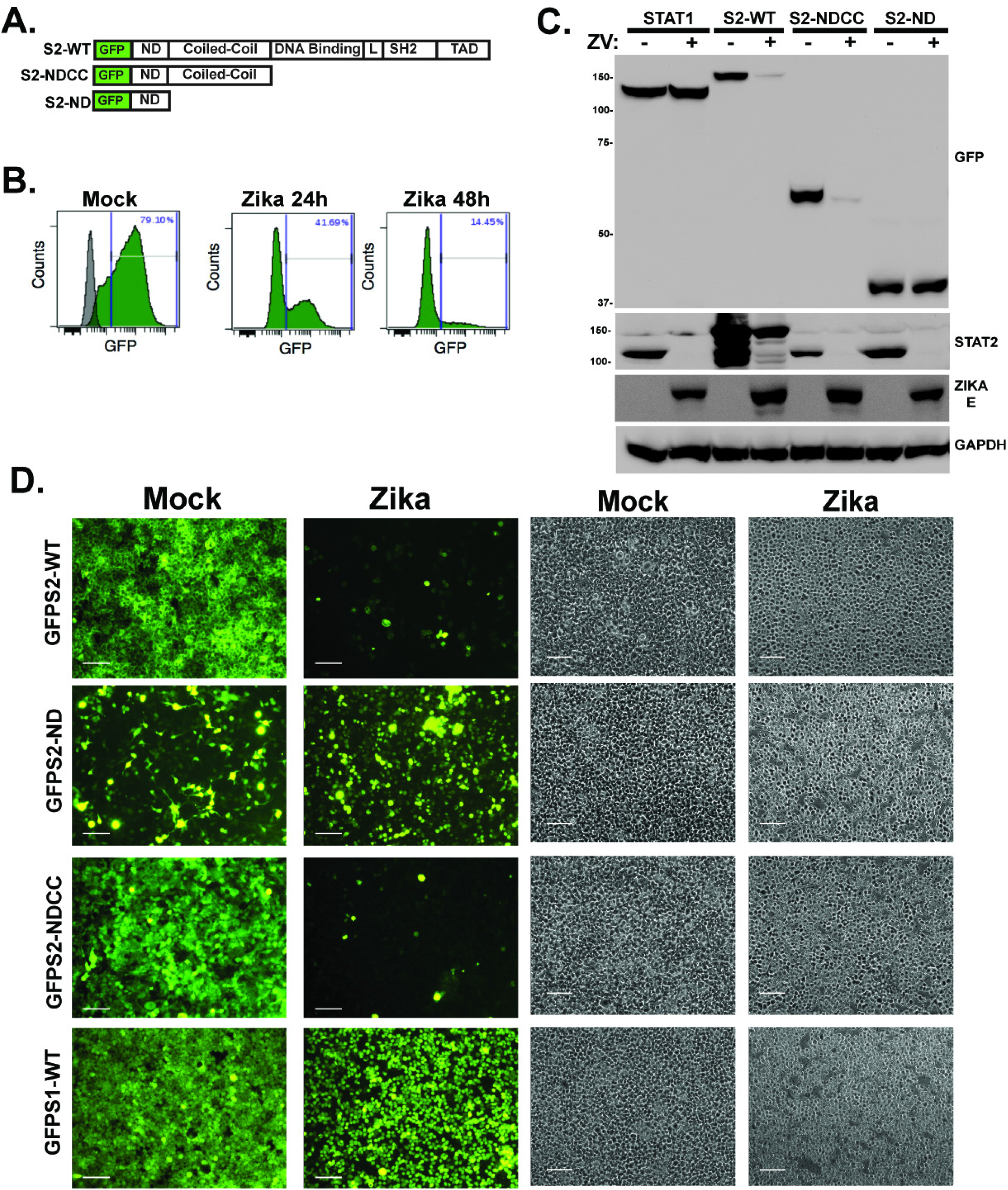
STAT2 coiled-coil domain is sufficient for Zika virus NS5 recognition and degradation. A.) Diagram of GFP-STAT2 fusion proteins carrying full length STAT2, NDCC (aa 1-315), or ND (aa 1-140). B.) Quantification of ZV -mediated GFP-STAT2 degradation. Flow cytometry was used to measure GFP fluorescence in mock-infected, or ZV -infected GFP-STAT2-WT expressing cells. Cells were infected with 5pfu/cell and analyzed at 24 and 48hpi. Grey peak overlay indicates the background fluorescence level of control HEK293 cells. C.) Cell lines expressing GFP fusions were infected with ZV and lysates prepared at 48hpi for immunoblotting with antiserum for GFP, STAT2, ZV E protein, or control GAPDH. D.) Cells expressing indicated GFP fusion proteins were imaged 48h after mock infection or ZV infection. GFP fusions with full-length STAT2 and STAT2-NDCC were efficiently degraded, but fusions with STAT2-ND or the negative control, STAT1, were unaffected by ZV infection. Bar=200 microns. Corresponding phase contrast images shown in parallel.

To directly observe the effects of ZV on STAT2 and its fragments, the GFP-STAT2 fusion proteins were visualized with fluorescence microscopy (Figure 3D). The subcellular distribution of GFP-STAT2 was found to reflect the known cytoplasmic distribution of endogenous STAT2, and fluorescence was dramatically eliminated by ZV infection. Unlike WT STAT2, the GFP-STAT1, GFP-STAT2-NDCC, and GFP-STAT2-ND were not retained in the cytoplasm and were more generally distributed in the cells. Infection with ZV resulted in efficient degradation of GFP-STAT2-NDCC, but GFP-STAT1 protein and GFP-STAT2-ND fluorescence remained intact after ZV infection. These findings indicate the NDCC is responsible for directing ZV-mediated degradation.

### STAT2 Coiled-Coil is essential and sufficient for ZV recognition and degradation

The inability of the ND to mediate GFP fusion protein degradation after ZV infection suggests that it may be dispensable for STAT2 targeting, and the key to STAT2 recognition and degradation may be encoded in the CC domain. To test this hypothesis, GFP fusions to the CC domain were constructed. Visualization of the GFP-STAT2-CC degradation by microscopy was confounded by an observed intrinsic instability of both N-terminal and C-terminal GFP fusion proteins, giving rise to low abundance of the desired GFP fusion proteins relative to GFP-NDCC (Figure 4A). The STAT2-CC-GFP fusion was more stable than GFP-STAT2-CC, and microscopic analysis of several independent clonally-selected stable lines support ZV-dependent degradation, despite their weak overall fluorescent signals (Figure 4 D, E). Despite this complication, immunoblotting was able to be used to evaluate both NS5 protein interaction and fusion protein degradation. Expression of STAT2-CC-GFP fusion protein along with FLAG-tagged NS5 resulted in specific co-precipitation of the STAT2-CC fusion (Figure 4B). In addition to NS5 interaction, the STAT2-CC fusion also was targeted for degradation after ZV infection (Figure 4C). These findings indicate that the STAT2 CC domain is an independent domain susceptible to recognition by NS5 and degradation during ZV infection.

**Figure 4.**
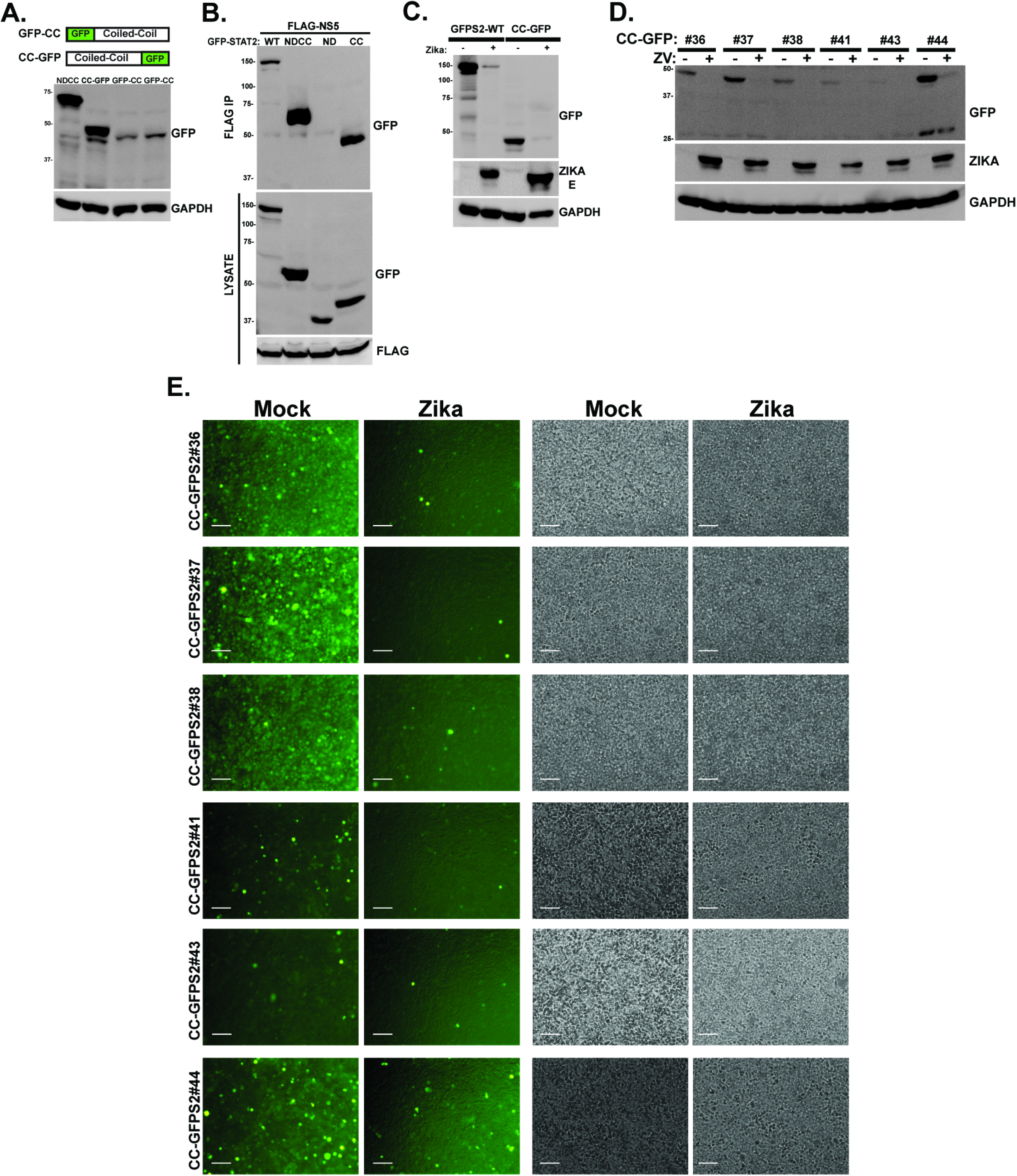
STAT2-CC domain is sufficient for recognition and degradation by ZV NS5. A.) *Top:* diagrams illustrate complementary GFP fusions with the STAT2 CC domain. *Bottom:* Immunoblot with GFP antiserum reveals relative steady state expression levels of STAT2-CC fusion proteins. B.) STAT2-CC-GFP is recognized by ZV NS5. Co-immunoprecipitation assays carried out as in Figure 2B, with GFP fusion proteins indicated. C.) STAT2-CC-GFP is degraded by ZV infection. Cells expressing GFP-STAT2 or STAT2-CC-GFP were mock infected or infected with 5pfu/cell ZV for 48h prior to immunoblotting with antisera for GFP, Zika E protein, or GAPDH. D.) Immunoblot of lysates from STAT2-CC-GFP clones in HEK293 cells before and after infection with 5 pfu/ml ZV for 48h. GFP antibody is used to detect STAT2-CC-GFP and ZV infection verified by E protein antiserum. E.) Fluorescence (left) and phase contrast (right) micrographs of STAT2-CC-GFP clones before and after infection with 5 pfu/ml ZV for 48h. Bar= 200 microns.

### STAT2 Coiled-Coil degradation by Zika virus requires the proteasome

Zika virus is thought to induce STAT2 degradation via the proteasome (24). To rule out alternate or off-target mechanisms for destabilization of the GFP-STAT2 fusion proteins and to validate the ability of the STAT2 CC domain to act as a discrete degron for degradation by the proteasome, the effects of proteasome inhibitors on ZV-induced degradation was evaluated. ZV infection efficiently eliminated endogenous STAT2 in control cells, and this was inhibited by both MG132 and epoxomicin treatment (Figure 5A). Similar experiments were conducted with cell lines expressing GFP-STAT2-WT, GFP-STAT2-NDCC or STAT2-CC-GFP (Figure 5B-D). All the GFP fusion proteins were efficiently degraded following ZV infection, but they were protected by treatment with proteasome inhibitors. These results demonstrate that the STAT2 CC domain functions as an independent degron that can target a heterologous protein (GFP) for proteasomal degradation during ZV infection.

**Figure 5.**
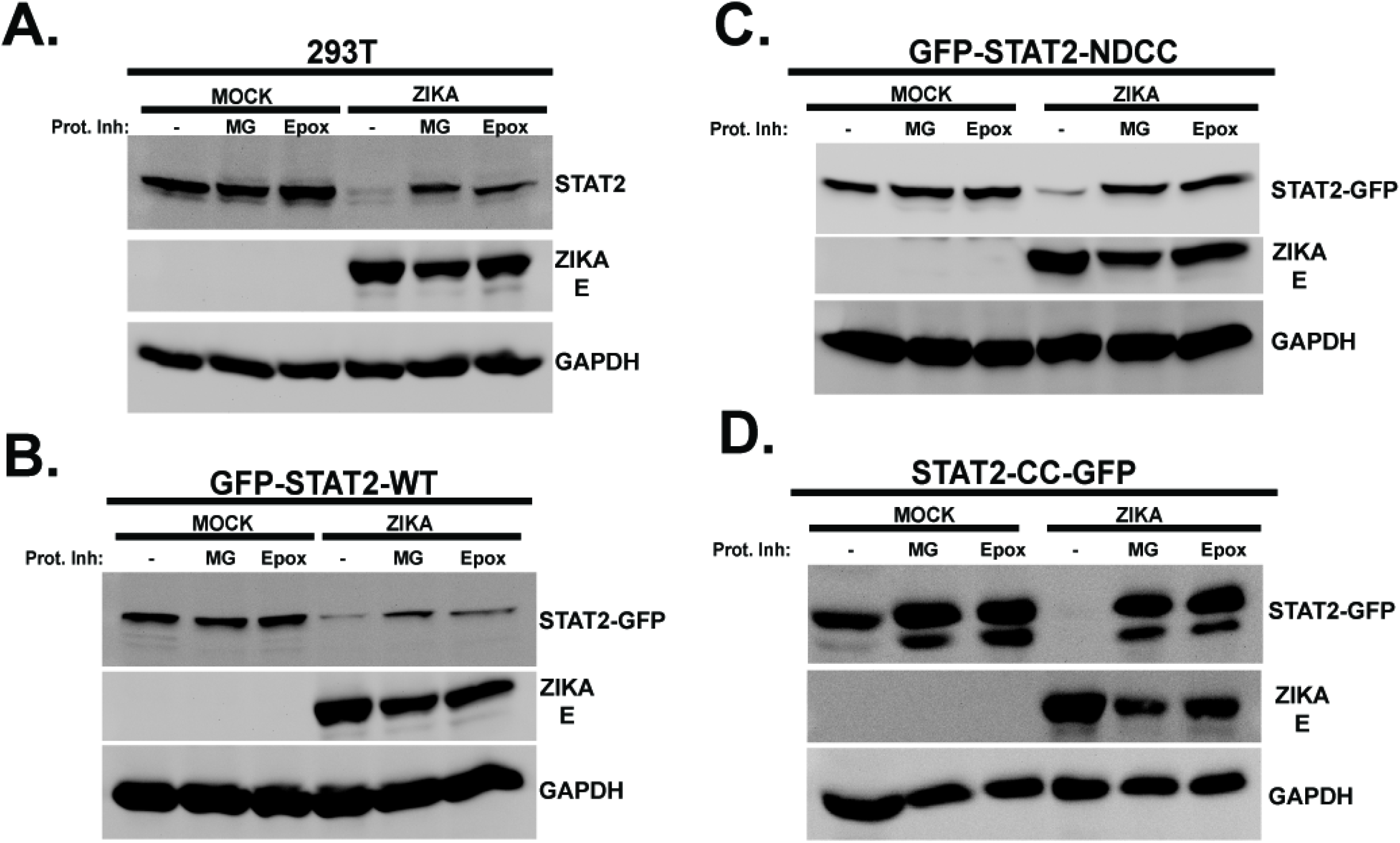
STAT2-CC domain degradation requires the proteasome. Control HEK293T cells (A.) or cells stably expressing GFP fusions with full-length (WT) STAT2 (B.), the STAT2-NDCC (C.), or the STAT2-CC (D.) indicated were mock infected or infected with ZV in the presence (+) or absence (-) of proteasome inhibitors, MG132 (MG) or Epoxomycin (Epox). Lysates were prepared and probed with antiserum for STAT2 C-terminus, GFP, Zika E protein, or GAPDH as indicated.

### Two helices of the STAT2 Coiled-coil mediate NS5 interaction and degradation

The STAT2 CC consists of four α-helices that present a predominantly hydrophilic surface area for interaction with other proteins in signal transduction and gene regulation (31, 32). To further dissect the STAT2 CC interaction and targeting by ZV NS5, cell lines stably expressing proteins encoding 3-helix (1-286), 2-helix (1-250), and 1-helix (1-187) fragments linked to GFP-ND were compared to GFP-STAT2-NDCC (1-315) which has all four helices (Figure 6A). The fragment containing STAT2 amino acids 1-250 (α1,2) retained the ability to co-precipitate with ZV NS5. However, further loss of 23 or more additional amino acids (1-227, 1-209, and 1-187) prevented NS5 co-precipitation, defining the first two helices of the STAT2 CC as crucial for ZV recognition (Figure 6B). To determine if this region can also mediate degradation, ZV was used to infect cells expressing GFP-STAT2-286 (α1,2,3) and GFP-STAT2-250 (α1,2). Consistent with their NS5 association, both of these proteins were degraded by ZV infection as determined by immunoblot (Figure 6C) as well as fluorescence microscopy (Figure 6D). These experiments implicate the first two α-helices of the STAT2 CC domain as the essential targeting region for recognition and degradation after ZV infection.

**Figure 6.**
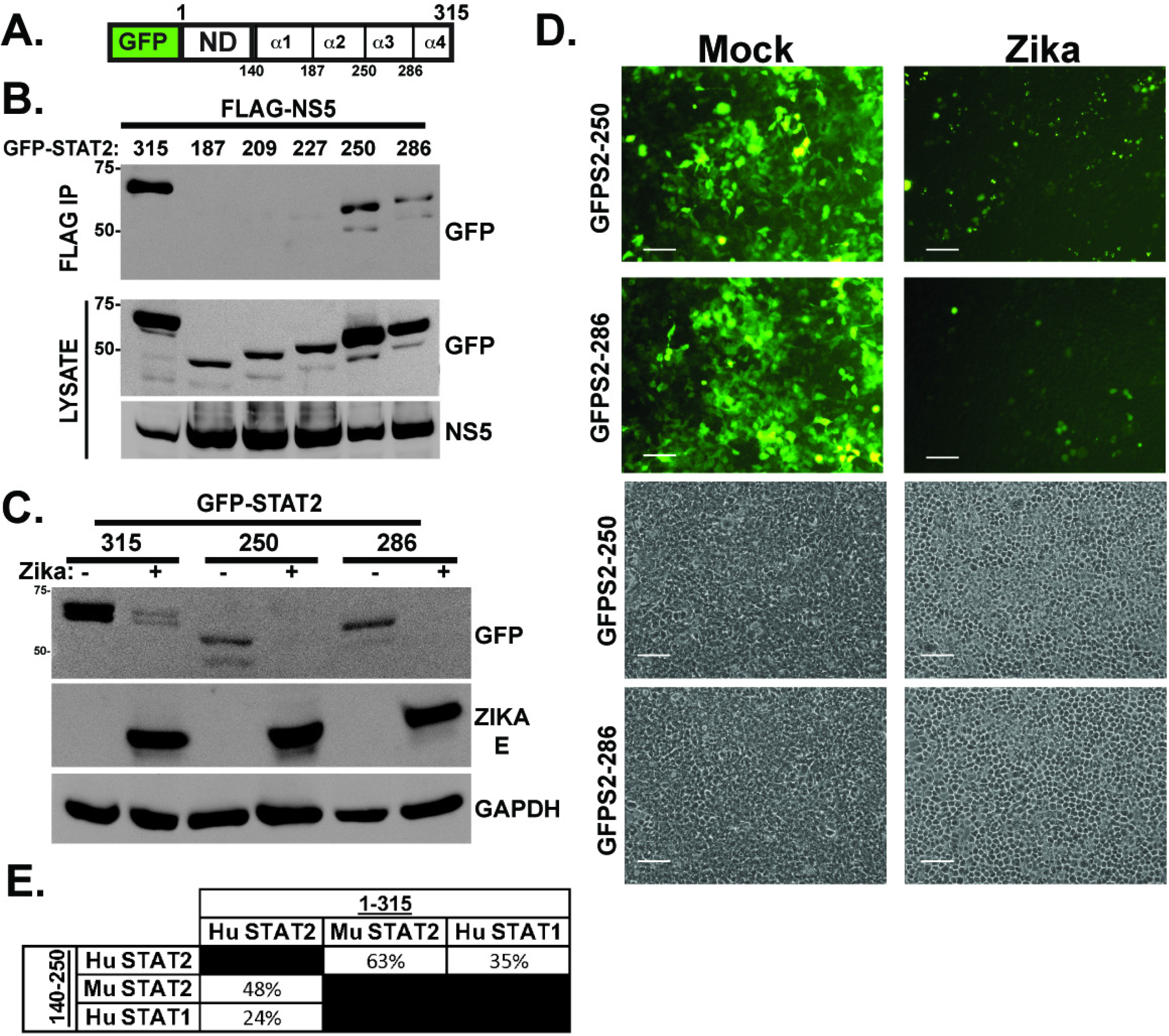
Zika virus degradation and NS5 interaction requires only the first two helices of the STAT2 CC domain. A.) Diagram of GFP-STAT2-NDCC indicating boundaries of 4 α-helices of the STAT2 CC domain. B.) Helices α3 and α4 are dispensable for NS5 interaction. Co-immunoprecipitation assay using the indicated GFP-STAT2-NDCC domain fragments along with FLAG-tagged NS5. C.) Zika virus induces degradation of GFP-STAT2 fragments 1-250 and 1-286. Immunoblot carried out as in Figure 3C, but using indicated GFP fusion proteins. D.) Zika virus induces degradation of GFP-STAT2 fragments 1-250 and 1-286; fluorescence microscopy carried out as in Figure 3D, but using cells expressing indicated GFP fusion proteins. Bar=200 microns. Corresponding phase contrast images shown below. E.) Comparison of the amino acid sequence identity among human (Hu) and mouse (Mu) STAT2 and human STAT1 in the NDCC (1-315) or the first two helices of the CC domain (140-250).

## Discussion

Virus replication in a host cell requires not only the fundamental processes of unpackaging, genome replication, mRNA transcription, and assembly of new particles, but also the passive or active suppression of host antiviral immune responses. For most RNA viruses, the IFN antiviral response system is a primary target of immune suppression, and the IFN-responsive STAT protein, STAT2, is a common target for flaviviruses including Zika, Dengue, and Yellow fever viruses (33).

STAT2 degradation is a conserved feature for these viruses, but the precise mechanisms and cellular partners needed for IFN signaling disruption are not well conserved between virus species. As a result, it is essential to functionally evaluate each viral mechanism systematically to determine their vulnerabilities and idiosyncrasies to inform their exploitation for therapeutic advantage. Zika virus can actively eliminate STAT2 to prevent IFN responses, using the viral non-structural protein, NS5, to target the typically stable STAT2 protein for efficient proteasome-mediated degradation (24). The experiments described here functionally dissect the interacting regions of ZV NS5 protein and its cellular target, STAT2. Data indicate that the STAT2 N-terminal 315 amino acids are key components for ZV NS5 recognition and degradation, corresponding with the STAT2 N-domain (ND) and coiled-coil (CC). Substitution of these 315 amino acids was sufficient to transfer the targeting ability of NS5 to a STAT2-STAT1 chimeric protein. This region is only ∼35% identical between STAT1 and STAT2 (Figure 6E), accounting for the specificity of NS5 interaction and degradation for STAT2. Expression of the STAT2 ND or CC domains as GFP fusions was used to delineate the minimal region(s) that are sufficient to act as an NS5 target. Experiments revealed that not only did GFP fused to the STAT2 NDCC domains bind to NS5 and rapidly turn over in infected cells, but that the CC domain alone (aa 140-315) was sufficient to mediate NS5 interaction and Zika-dependent degradation via the cellular proteasome.

Sequential elimination of the four α-helices of the STAT2 CC indicate that helix 1 and helix 2 are essential for NS5 interaction and degradation and, but helix 3 and helix 4 are dispensable, narrowing the essential region for determining STAT2 targeting specificity to amino acids 140-250. STAT2 targeting by ZV is known to be species-restricted; the murine STAT2 protein is not degraded in infected cells and does not interact with NS5 (24). Alignment of the human and mouse STAT2 amino acid sequence in the NDCC region (1-315) indicates it is 63% identical, but focusing on amino acids 140-250 reveals only 48% sequence identity (Figure 6E). This lower sequence conservation may account for species-specific STAT2 targeting by ZV NS5. These sequence differences corresponding to differential ZV NS5 targeting support a role for the identified degron as a crucial viral target for highly specific host IFN evasion, and therefore represents a novel therapeutic target.

During the course of these investigations, crystallographic and cryo-electron microscopy structures of the Zika virus and Dengue virus NS5-STAT2 complex were described that provide further support of our functional analysis (30). The structures illustrate the importance of the CC domain in forming an NS5-STAT2 complex. The two domains of NS5, the RNA-dependent RNA polymerase (RDRP) and the methyltransferase (MT), were both implicated in STAT2 contact. The NS5 RDRP fragment was found to associate with helices 1 and 2, and conserved STAT2 residues F175 and R176 established key contacts with NS5. Atomic models based on the cryo-EM structures show that the STAT2 CC is anchored at an interdomain cleft formed between the NS5 MT and RDRP domains, indicating that both domains of NS5 interact with STAT2. These results strongly support our conclusion that the primary functional ZV target site is the STAT2 CC domain and invoke the importance of amino acids 140-250. Interestingly, the same region of the STAT2 CC is required for IRF9 binding to form the IFN-activated ISGF3 transcription complex heterotrimer (32). Functional antagonism between hSTAT2 and ZIKV NS5 in prevention of ISGF3 assembly was observed in competition assays, indicating multi-level NS5 IFN suppression are determined by the STAT2 CC interaction (30).

The structures also recognized a contact site between the STAT2 ND and the NS5 RDRP fragment, mediated by an invisible linker between the ND and the CC. As our results strongly indicate that the STAT2 ND itself does not co-precipitate with NS5 or undergo degradation during ZV infection, the observed ND-RDRP interaction may represent a low-affinity interaction that could participate in other aspects of NS5 activity. Alternatively, the ND engagement of NS5 may reflect a means by which STAT2 counter-antagonizes NS5 polymerase action as an antiviral measure.

In summary, the results presented here define a transferable NS5-mediated targeting sequence that allows ZV to overcome IFN antiviral immunity by proteasomal degradation of STAT2. As a key feature of ZV immune escape, this region may be exploited as a target for development of antiviral compounds or other therapeutic strategies.

## Materials and Methods

### Cells, Viruses, and Inhibitors

HEK293, Bosc, 293T, CV1, HeLa, 2fTGH and Vero cells were grown in DMEM (Invitrogen, cat# 11965118) supplemented with 10% cosmic calf serum (CCS, Hyclone, cat# 25200114) and 1% penicillin-streptomycin (Invitrogen, cat# 15140122). Cells are routinely tested for mycoplasma contamination and regularly restored from early passage frozen stocks. Zika virus (ATCC, cat# VR-84, MR 766 strain) was propagated and titrated in Vero cells. Infection of cells with Zika virus was performed at an m.o.i of 5 pfu/cell in serum-free media. After 1h, cells were washed, placed in growth medium supplemented with 2% CCS, and harvested 48h later. Proteasome inhibitors MG132 (20μM; Selleck Chemicals, cat# S2619) and Epoxomycin (400nM; R&D Systems, cat# I-110) were incubated with cells for 12h. Stable cell lines were generated by expressing GFP-STAT2 plasmids in HEK293 cells and selecting with 500μg/mL G418. Positive clones were ultimately screened by immunoblot with a GFP antibody and high-expressing cells were selected by fluorescence activated cell sorting.

### Plasmids and mutagenesis

To generate HA-tagged NS5, Vero cells were infected with Zika virus MR 766 strain, total RNA was isolated using Trizol (Invitrogen, cat# 15596018), and cDNA was amplified by PCR with gene-specific primers and cloned into pcDNA3-HA expression vector. pLV_Zika_NS5_Flag was obtained from Vaithi Arumugaswami (Addgene plasmid # 79639); and used to transfect cells directly for FLAG protein expression.

STAT2 expression vectors, fragments, and STAT1 chimeras were previously described (34-36). To generate GFP fusion proteins, full length STAT2 ORF was amplified by PCR and subcloned into mammalian expression plasmid (pEGFP-C1). Truncations were generated by stop codon insertion mutagenesis with QuikChange II XL mutagenesis kit (Agilent, cat# 200522), and GFP fusion expressing the STAT2 CC domain alone was constructed by PCR with specific primers to enable cloning into plasmid pEGFP-N1 or pEGFP-C1. All constructs confirmed by DNA sequencing.

### Immunoprecipitations and immunoblotting

For co-immunoprecipitation experiments, FLAG-tagged and HA-tagged or GFP-tagged plasmids were transfected in HEK293T cells by the calcium phosphate method. 48h post transfection, cells were harvested by first washing with cold phosphate-buffered saline and then lysed with whole cell extract buffer (WCEB) consisting of 50mM Tris, 280 mM NaCl, 0.5% NP-40, 0.2mM EDTA, 2mM EGTA, 10% glycerol, 1mM DTT, 2.5mM sodium vanadate, and protease inhibitors. Cell lysates were then precleared with sepharose beads. A percentage of the cleared lysates was reserved for analysis and the remainder incubated with FLAG M2 affinity beads (Sigma, cat# A2220) overnight and washed 5X with WCEB. FLAG and HA IPs were eluted with SDS sample buffer, FLAG and GFP IPs were eluted with 3XFLAG Peptide (ApexBio, cat# A6001), then separated by SDS-PAGE and processed for immunoblotting. For immunoblotting, the separated proteins were transferred to nitrocellulose and probed with commercial primary antibodies recognizing FLAG (Sigma, cat# F3165), HA (Sigma, cat# H3663), GFP (Santa Cruz Biotechnology, cat# sc-9996), STAT2-C (Santa Cruz Biotechnology, cat# sc-476), STAT1-C (Santa Cruz Biotechnology, cat# sc-345), ZIKA E (Millipore Sigma, cat# MABF2046), or GAPDH (Santa Cruz Biotechnology, cat# sc-47724). Proteins were visualized by enhanced chemiluminescence (PerkinElmer, cat# NEL105001EA). Figures show representative images of 3-5 biological replicates.

### Reporter gene assays

HEK293 and 2fTGH cells were transfected with a 5xISRE luciferase reporter gene along with a *Renilla* luciferase vector and expression vectors for Zika NS5, PIV5 V and HPIV2 V. 24 hours post-transfection, cells were stimulated for 8h with IFNα (1000 units/mL) then harvested and assayed for firefly and *Renilla* luciferase activities using the Dual Luciferase reporter assay system (Promega, cat# E1960). Relative luciferase activity was calculated by dividing the firefly luciferase values by those of the *Renilla* luciferase. Data are plotted as mean values, with error bars representing standard deviation.

### Fluorescence Microscopy and Imaging

Cell Sorting was performed at the Northwestern Flow Cytometry Core Facility on a BD FACSAria SORP system and BD FACSymphony S6 SORP system, purchased through the support of NIH 1S10OD011996-01 and 1S10OD026814-01. Fluorescent images were obtained using a Nikon Eclipse Ti fluorescence microscope equipped with a Photometrics CoolSnap CF2 camera at 10X magnification. Images were adjusted for brightness and contrast using Adobe Photoshop.

## Acknowledgements

Supported by NIH grant R21 AI148949-01 to CMH. JJL and GA were supported by Cellular and Molecular Basis of Disease Training Grant (NIH T32 GM008061), and the Northwestern University Flow Cytometry Core Facility is supported by Lurie Cancer Center support grant NCI CA060553, NIH 1S10OD011996-01 and 1S10OD026814-01.

